# First haplotype-resolved genome assembly of citral-rich lemongrass *Cymbopogon flexuosus* var. Krishna

**DOI:** 10.64898/2026.02.17.706310

**Authors:** Swati Tyagi, Vikrant Gupta, Sanjeet Verma, Neelam Prabha Negi, Sanjay Kumar, Prabodh Kumer Trivedi

## Abstract

*Cymbopogon flexuosus* var. Krishna (lemongrass) is an aromatic grass valued for its high citral content, which is widely used in the fragrance, flavor, and pharmaceutical industries. *C. flexuosus*, a member of the Poaceae family, is a predominantly outcrossing species characterized by a highly heterozygous genome. Despite its economic importance and widespread cultivation, a high-quality reference genome has been lacking. Here, we report the first chromosome-scale genome assembly of lemongrass, generated using PacBio HiFi long-read sequencing combined with Omni-C chromatin conformation capture data. The resulting pseudo-haploid assembly spans approximately 798 Mb, organized into 10 chromosomes, and exhibits a scaffold N50 of 64.35 Mb. The assembly demonstrates high completeness, with 99.8% BUSCO recovery, and comprises ∼37,254 predicted protein-coding genes. In addition, we generated haplotype-resolved assemblies that capture the allelic diversity of this heterozygous genome. The haplotypes have sizes of ∼750 Mb and ∼726 Mb, representing 95–98% of the pseudo-haploid genome, and together they provide phase-resolved information for gene families and biosynthetic pathways. These high-quality assemblies establish a foundational genomic resource for advancing molecular breeding, comparative genomics, and metabolic engineering of lemongrass and related aromatic grasses.

## Background & Summary

*Cymbopogon flexuosus* (East Indian lemongrass) is one of the most important aromatic grasses of the Poaceae family, valued globally for its essential oils rich in citral, a precursor of vitamin A, ionones, and various aroma compounds used in flavour, fragrance, cosmetics, and pharmaceuticals ^1^. Beyond its industrial uses in flavours, fragrances, and cosmetics, lemongrass also holds significance in traditional medicine owing to antimicrobial, anti-inflammatory, and anxiolytic properties ^2-4^. India is one of the leading cultivators and exporters, with ∼20,000 hectares under cultivation, providing sustainable income to small and marginal farmers, particularly on less fertile and underutilized lands ^5^. The lemongrass species exhibits significant morphological and chemo-typical diversity, with natural populations comprising different ploidy levels including diploid, tetraploid, and hexaploid races ^6-8^. Despite its commercial importance, genomic resources for *C. flexuosus* remain scarce compared to major cereals such as rice (*Oryza sativa*) ^9^, maize (*Zea mays*) ^10^, and sorghum (*Sorghum bicolor*) ^11^, where high-quality genomes have transformed functional biology and crop improvement. For *Cymbopogon*, existing studies have been limited to transcriptomes ^1,12,13^ or draft assemblies of related species such as *C. citratus* ^*14*^, which do not provide the resolution necessary for understanding genome organization, secondary metabolite gene clusters, or for enabling genomics-assisted breeding. Among cultivated varieties, *C. flexuosus* var. Krishna (Fig. 1a), released by CSIR-Central Institute of Medicinal and Aromatic Plants (CIMAP), is recognized as an elite cultivar with high citral content (78–82 percent), stable yield, and adaptability across diverse agro-climatic regions of India ^*5,15*^. It is the most widely cultivated lemongrass variety in northern and central India and a primary contributor to India’s essential oil exports ^16^. Given its superior essential oil profile and economic importance, Krishna represents a strategic target for genomic resource developments ^16-18^.

**Fig 1.**
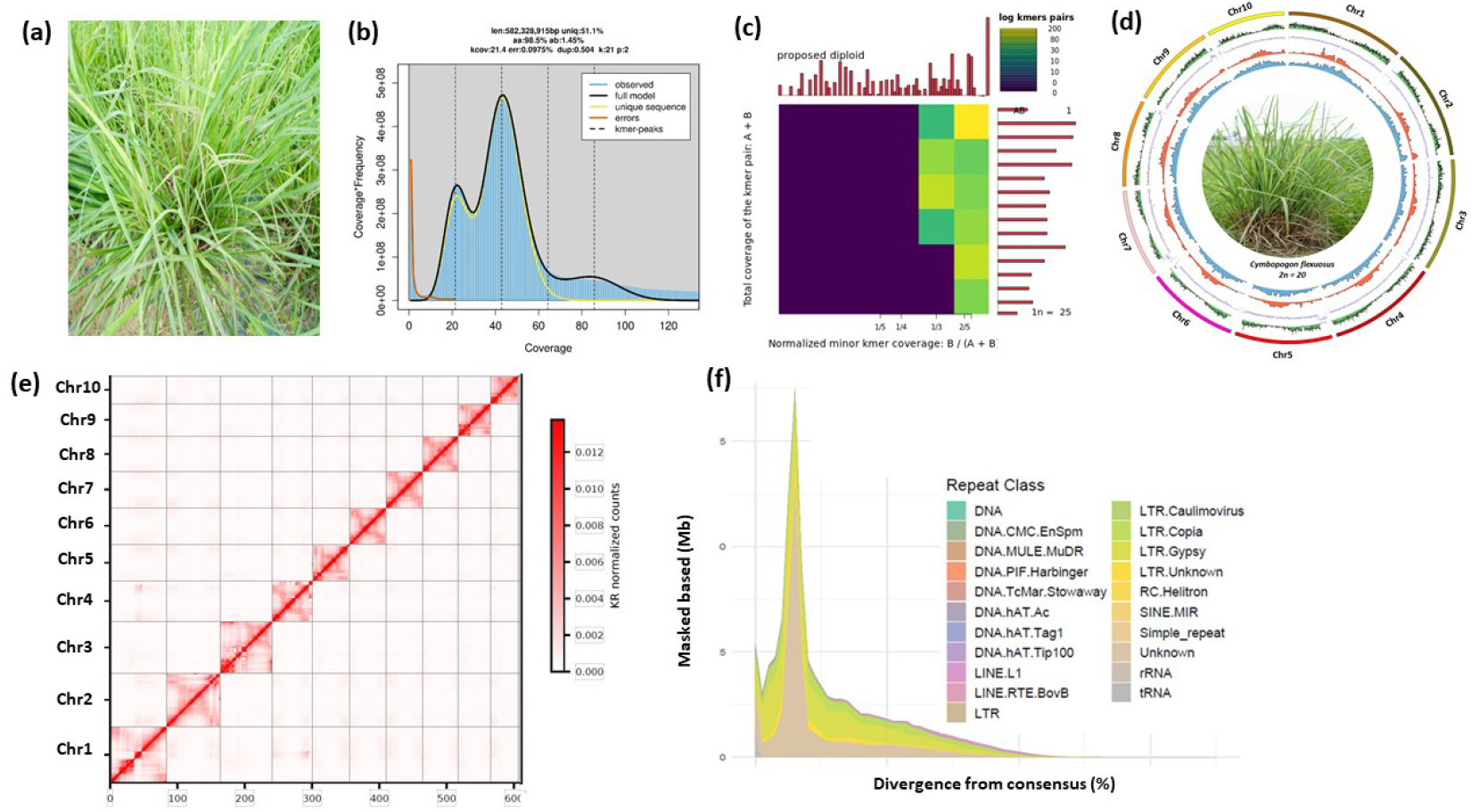
Genome sequencing and assembly overview for *Cymbopogon flexuosus* (lemongrass). (a) Representative photograph of *C. flexuosus* var. Krishna plant. (b) K-mer frequency histogram from Illumina short-read data, showing major peaks for heterozygous and homozygous k-mers. The plot estimates genome size, heterozygosity, and repeat content prior to assembly. (c) smudgplot analysis heatmap and summary plot illustrating normalized minor k-mer coverage, supporting a diploid genome structure. Data aggregation shows the majority of k-mer pairs clustering as expected for diploid plants. (d) Circular visualization of the chromosome-level genome assembly with ten pseudo-chromosomes, displaying gene density, repeat content, and key genomic features. Outer to inner ring: pseudo-chromosome, GC content, Repeat/Gypsy, Repeat/Copia and at the center shows a lemongrass plant; (e) Hi-C contact map validating chromosome-scale assembly and contiguity, with strong signals along the diagonal indicating well-resolved chromosome structures. (f) Distribution and classification of repetitive elements annotated in the assembled genome. The plot shows repeat class breakdown (DNA transposons, LTR retrotransposons, etc.) and their divergence from the consensus, with masking statistics in megabases.

In this study, we successfully generated a high-quality genome assembly for *Cymbopogon flexuosus* var. Krishna. Using PacBio HiFi and Omni-C sequencing, we produced both a pseudo-haploid assembly and haplotype-resolved sequences, resulting in the first chromosome-level reference genome for lemongrass. This resource represents a major step forward and provides a strong foundation for future research on this important crop.

## Methods

### Plant material and sample preparation

Young, healthy leaves of *C. flexuosus* var. *Krishna* from a single plant were obtained from CSIR-Central Institute for Medicinal and Aromatic Plants, Lucknow, India. The plant was maintained under controlled greenhouse conditions at tissue culture and transformation facility (TTF), CSIR-CIMAP following the standard protocols. Fresh tissue was harvested in the early morning to minimize polysaccharide accumulation. High molecular weight genomic DNA was extracted using the Qiagen MagAttract HMW DNA kit following the manufacturer’s protocol with modifications to reduce polysaccharide and secondary metabolite contamination. DNA quality was assessed using pulsed-field gel electrophoresis and a Femto Pulse system, while concentration was quantified with Qubit fluorometry and Nanodrop spectrophotometry. DNA samples with a mean fragment size >50 kb and A260/280 ratio between 1.8–2.0 were used for library preparation as discussed. ^19^

### HiFi and Omni-C library preparation and sequencing

The genome size of lemongrass was estimated from PacBio HiFi reads (Table 1) using a 21-mer analysis with Jellyfish. A k-mer frequency distribution was generated and further analyzed with GenomeScope 2.0, which provided estimates of genome size and heterozygosity. The estimated genome size was in between 0.58-0.80□Gb (based on kmer and flow cytometer analysis) with heterozygosity rate of 0.8% and repetitive rate of ∼48.9% (Fig. 1b). In addition, smudgeplot analysis confirmed the diploid nature of the genome (Fig. 1c). SMRTbell libraries were prepared using the SMRTbell Express Template Prep Kit 3.0 (PacBio), optimized for long insert libraries. DNA repair and end-polishing steps were performed following the manufacturer’s protocol. Libraries were size-selected using a BluePippin system (Sage Science) with a cutoff of ∼15–20 kb. Sequencing was performed on the PacBio Revio platform using Revio SMRT Cells, generating high-fidelity (HiFi) reads with the latest chemistry. CCS (circular consensus sequencing) reads were produced with the PacBio CCS v6.4 pipeline, requiring a minimum of three passes per molecule and a predicted accuracy ≥0.99. In total, ∼32 Gb of HiFi reads (∼40× coverage of the estimated genome size) were generated (Table 1). For scaffolding, an Omni-C library was prepared using the Dovetail Omni-C Kit (Dovetail Genomics) following the manufacturer’s instructions. Crosslinked chromatin was digested and proximity-ligated to generate chimeric junctions representing physical contacts across the genome. The library was sequenced on an Illumina NovaSeq 6000 platform to produce 150 bp paired-end reads. Approximately 377 million read pairs (∼150× coverage) were generated (Table 2).

**Table 1.** General Statistics of PacBio HiFi sequencing data used in ‘Krishna’ genome assembly.

| Metrics | Value |
| --- | --- |
| Number of reads | 3525146 |
| Total bases (bp) | 31395198794 |
| Minimum length of reads (bp) | 91 |
| Maximum length of reads (bp) | 45479 |
| Average length of reads (bp) | 8906.07 |

**Table 2.** Summary statistics of Omni-C sequencing data used in ‘Krishna’ genome assembly.

| Metrics | Value |
| --- | --- |
| Number of read pairs | 377444682 |
| Average length of reads (bp) | 159 |
| Total bases (bp) | 1.20027E+11 |
| GC (%) | 52.87 |

### Pseudo-haploid Genome assembly

The pseudo-haploid genome assembly of ‘Krishna’ genome was generated using Hifiasm (v.0.16.1) with default parameters optimized for HiFi reads ^20^ (Fig. 1d, Table 3). Redundant haplotigs were purged using Purge_dups (v.1.2.5) ^21^ to produce a haploid representation of the genome. Scaffolding was performed with HapHiC (v. 1.0.6) using Omni-C data, anchoring contigs into chromosome-scale scaffolds. The draft assembly comprised 623 scaffolds, totalling 683,312,799 bp, with an N50 of 42.25 Mb. These scaffolds were anchored onto 10 chromosomes, yielding a cumulative length of 1,135,979,557 bp (Table 3, Fig. 1d). The resulting chromosome-level assembly spanned 798 Mb, with a scaffold N50 of 64.35 Mb, and the 10 pseudo-chromosomes together accounted for ∼98% of the genome. Assembly completeness was evaluated using BUSCO (v5.4.3) against the embryophyta, eukaryota, and poales lineage datasets, which confirmed high genome integrity with 99.98%, 98.0%, and 95.4% completeness, respectively (Table 3). Gap filling was performed using Minimap2 (v2.26-r1175)^22^ and LR-GapCloser (v1.2.1)^23^ reducing the assembly to only 38 remaining gaps (Table 4). Structural accuracy was evaluated using Merqury v1.3 which provided k-mer–based quality value (QV) estimates and base-level completeness metrics ^24^. Hi-C contact maps, visualized in Juicebox v1.11.08, confirmed the accuracy of chromosome anchoring (Fig. 1e). Haplotype-resolved assemblies were further generated from the unitig assembly obtained through *hifiasm*, following the approach described by Mascher *et al*. (19). The final manually curated pseudomolecules for the two haplotypes spanned 750 Mb (Hap1) and 726 Mb (Hap2), respectively (Table 3). Chromosomes within each haplotype were designated according to their corresponding chromosomes in the pseudo-haploid reference. In total, 10 chromosomes were grouped for Hap1 and Hap2, providing a fully resolved, chromosome-level haplotype assembly (Table 3).

**Table 3.** Summary statistics of the ‘Krishna’ genome assembly.

| Assembly characteristics | Hap1 | Hap2 | Psuedohaploid |
| --- | --- | --- | --- |
| Number of contigs | 883 | 382 | 673 |
| Contig size (bp) | 7.50E+08 | 7.25E+08 | 798944333 |
| Contig N50 (bp) | 10891133 | 16705924 | 52680464 |
| Number of scaffolds | 762 | 267 | 635 |
| Scaffold size (bp) | 7.50E+08 | 7.26E+08 | 798948133 |
| Scaffold N50 (bp) | 53085257 | 58722762 | 64355099 |
| Scaffold N90 (bp) | 3327303 |  | 3327303 |
| Number of chromosomes | 10 | 10 | 10 |
| Lineage (Eukaryota) |  |  |  |
| Complete BUSCOs (Number / %) | 424 / 99.8% | 424 / 99.8% | 424 / 99.8% |
| Complete and single-copy BUSCOs (Number / %) | 405 / 95.3% | 405 / 95.3% | 405 / 95.3% |
| Complete and duplicated BUSCOs (Number / %) | 19 / 4.5% | 19 / 4.5% | 19 / 4.5% |
| Fragmented BUSCOs (Number / %) | 1 / 0.2% | 1 / 0.2% | 1 / 0.2% |
| **Missing BUSCOs (Number / %)** | 0 / 0% | 0 / 0% | 0 / 0% |
| **Lineage (Embryophyta)** |  |  |  |
| **Complete BUSCOs (Number / %)** | 1582 / 98.0 % | 1583 / 98.1 % | 1581 / 98.0 % |
| **Complete and single-copy BUSCOs (Number / %)** | 1503 / 93.1 % | 1506 / 93.3 % | 1517 / 94.0 % |
| **Complete and duplicated BUSCOs (Number / %)** | 79.0 / 4.9% | 77.0 / 4.8 % | 64 / 4.0 % |
| **Fragmented BUSCOs (Number / %)** | 8 / 0.5 % | 11 / 0.7 % | 8 / 0.5 % |
| **Missing BUSCOs (Number / %)** | 24.0 / 1.5 % | 20.0 / 1.2 % | 25 / 1.5 % |
| **Total busco searched (Number / %)** | 1614 / 100 % | 1614 / 100 % | 1614 / 100 % |
| **Lineage (Poales)** |  |  |  |
| **Complete BUSCOs (Number / %)** | 4788 / 97.8% | 4819 / 98.4 % | 4669 / 95.4 % |
| **Complete and single-copy BUSCOs (Number / %)** | 4587 / 93.7 % | 4623 / 94.4 % | 4536 / 92.6 % |
| **Complete and duplicated BUSCOs (Number / %)** | 201 / 4.1 % | 196 / 4.0 % | 133 / 2.7 % |
| **Fragmented BUSCOs (Number / %)** | 35 / 0.7 % | 30 / 0.6 % | 61 / 1.2 % |
| **Missing BUSCOs (Number / %)** | 73 / 1.5 % | 47 / 1.0 % | 166 / 3.4 % |
| **Total busco searched (Number / %)** | 4896 / 100% | 4896 / 100% | 4896 / 100 % |

**Table 4.** Summary of the chromosome-level genome assembly of ‘Krishna’.

| **Chromosome number** | **Primary** |  |  |
| --- | --- | --- | --- |
|  | **Length (bp)** | **Gapcount** | **Gaplocus** |
| **Chr1** | 82045033 | 4 | 74658440-74658539, 74677664-74677763, 81407107-81407206, 81992614-81992713 |
| **Chr2** | 77913746 | 1 | 71707565-71707664 |
| **Chr3** | 78452326 | 10 | 76408775-76408874, 76948269-76948368, 77008108-77008207, 77018614-77018713, 78200655-78200754, 78229961-78230060, 78229961-78230060, 78263684-78263783, 78290654-78290753, 78430843-78430942 |
| **Chr4** | 76454182 | 3 | 101910-102009, 54582728-54582827, 76388597-76388696 |
| **Chr5** | 67983055 | 1 | 15785859-15785958 |
| **Chr6** | 64355099 | 3 | 60067898-60067997, 60673975-60674074,<br>64156990-64157089 |
| **Chr7** | 61062956 | 5 | 571597-571696, 611122-611221, 982015-982114, 2094299-2094398, 2281177-2281276 |
| **Chr8** | 60628767 | 5 | 21875308-21875407, 48214262-48214361,<br>48256277-48256376, 48271970-48272069,<br>60446749-60446848 |
| **Chr9** | 59840558 | 2 | 47600601-47600700, 59776347-59776446 |
| **Chr10** | 54577077 | 4 | 52680465-52680564, 52728943-52729042,<br>53506733-53506832, 54531892-54531991 |

### Pseudo-haploid assembly repeat annotation

Transposable elements (TEs) in the *C. flexuosus* var. *Krishna* genome was identified and annotated using an integrated pipeline that combined TRF (v.4.10), RECON, RepeatScout (v. 1.0.7), RepeatMasker (v4.1. 9), RepeatAfterMe (v.0.0.7)^25,26^. *De novo* repeat libraries were constructed with RepeatModeler (v2.0.7) ^27^ and EDTA (v2.2.0) ^28^ to improve classification accuracy. In total, 51,743,703 interspersed repeat elements were annotated, collectively representing 65.16% of the genome. These comprised both retrotransposons—long terminal repeat (LTR) and non-LTR elements—and DNA transposons with terminal inverted repeats (TIR) and non-TIR structures (Table 5, Fig. 1f). Among these, LTR-Gypsy retrotransposons were the predominant class, accounting for 26.41% of the assembled genome (Table 5). Detailed statistics, including counts, total lengths, and proportional contributions of each TE category, are summarized in Table 5.

**Table 5.** Overview of repetitive sequence composition in the ‘Krishna’ genome.

| Type |  | Number of elements |  | Length occupied | Percentage of genome |
| --- | --- | --- | --- | --- | --- |
| Retroelements | SINEs | 694 |  | 81265 bp | 0.01% |
|  | LINEs | RTE/Bov-B | 1975 | 1239607 bp | 0.16% |
|  |  | L1/CIN4 | 24282 | 16084527 bp | 2.01% |
|  | LTR | BEL/PAO | 87 | 42181 bp | 0.01% |
|  |  | Tyl/Copia | 31808 | 57781336 bp | 7.23% |
|  |  | Gypsy/DIRS1 | 82189 | 206835308 bp | 25.89% |
| DNA Transposons |  | hobo-Activator | 3589 | 1482941 bp | 0.19% |
|  |  | MULE-MuDR | 5895 | 3266099 bp | 0.41% |
|  |  | Tourist/Harbinger | 2929 | 968890 bp | 0.12% |
| Total interspersed repeats |  |  |  | 504374265 bp | 63.14% |

### Pseudo-haploid assembly gene prediction and annotation

Gene prediction in the *C. flexuosus* var. *Krishna* genome was performed using the MAKER pipeline (v3.01.03)^29^ which integrates ab initio predictions, protein homology, and transcriptomic evidence. Ab initio gene models were generated with Augustus (v3.4.0) ^30^, trained with *S. bicolor*. For transcriptome-based support, RNA-seq datasets derived from *C. flexuosus* leaves and inflorescences were aligned with HISAT2 (v2.2.1)^31^ nd assembled using StringTie (v2.1.7)^32^. Homology-based annotations incorporated protein sequences from *Oryza sativa, Sorghum bicolor, Zea mays*, and *Arabidopsis thaliana*. The integrated pipeline predicted a total of 37,254 protein-coding genes.

Functional annotation was carried out using a combination of approaches. Predicted proteins were searched against SwissProt/UniProt, NCBI nr, and Pfam (v35.0) using BLASTP and HMMER (v3.3.2). Functional domains were further characterized with InterProScan (v5.56-89.0) ^33^, and and Gene Ontology (GO) terms were assigned using Blast2GO (v6.0). Orthology-based annotation with eggNOG-mapper (v5.0)^34^ provided additional functional assignments, while the KEGG Automatic Annotation Server^35^ enabled mapping of genes to metabolic and signalling pathways. In total, 32,428, 31,516, 14,054, and 34,681 genes were annotated through UniProt, eggNOG-mapper, Pfam, and InterPro databases, respectively (Table 6). This process resulted in the annotation of 32,428, 31,516, 14054, and 34681 genes using Uniprot, eggNOG-mapper, Pfam and InterPro database, respectively (Table 6). Furthermore, eggNOG-mapper identified 24,784 genes with GO terms and 26,173 genes with KEGG Orthology KO) terms (Table 6). Genes involved in monoterpene biosynthesis pathways were specifically identified and manually curated based on homology to characterized terpene synthases.

**Table 6.** Summary of genome-wide annotation of functional genes.

| Metrics | Gene count |
| --- | --- |
| Protein coding gene | 37,254 |
| eggNOG-mapper | 31,516, |
| PANTHER | 32,428 |
| Pfam | 14,054 |
| InterPro | 34,681 |
| GO (eggNOG-mapper) | 24,784 |
| KEGG (eggNOG-mapper) | 26,173 |

### Data Records

The complete genome assembly and associated datasets have been deposited in public repositories with the given accessions. The final chromosome-level genome assembly of *C. flexuosus* var. *Krishna* has been deposited at NCBI GenBank under BioProject accession PRJNA1301599 with Assembly accession [JBQKUF000000000] and BioSample (SAMN50449607). The assembly consists of 10 chromosomes with total length of 0.79 Gb and contig N50 of 64.35 Mb.

### Technical Validation

The completeness of the gene space in the *C. flexuosus* var. *Krishna* genome assembly was evaluated using BUSCO (v5.4.6) (36) with the *embryophyta_odb10* dataset (n = 1,614). The analysis recovered 98.0% complete genes, comprising 94.0% single-copy and 4.0% duplicated, with 0.5% fragmented and 1.5% missing genes. These results are on par with other high-quality grass genome assemblies, underscoring the comprehensive recovery of the coding repertoire.

## Data Availability

The complete genome assembly and associated datasets have been deposited in public repositories with the given accessions. The genome sequencing data i.e. the HiFi sequencing data, the Hi-C sequencing data, and the RNA sequencing data were deposited in our in-house medicinal and aromatic plants database i.e. GR-MAP database. The final chromosome-level genome assembly of *C. flexuosus* var. *Krishna* has been deposited at NCBI GenBank under BioProject accession PRJNA1301599 with Assembly accession [JBQKUF000000000] and BioSample (SAMN50449607). The assembly consists of 10 chromosomes with total length of 0.79 Gb and contig N50 of 64.35 Mb.

## Code Availability

All computational analyses were performed using publicly available software packages with version numbers and parameters specified in the Methods section. Custom scripts for data processing, analysis pipelines, and visualization were developed using Python 3.8, R 4.2.0, and Bash scripting. No proprietary software was used in this study. All genome assembly and annotation pipelines can be reproduced using the provided documentation and publicly available computational resources. Specific software versions and database versions used are documented in the GitHub repository for complete reproducibility.

## Acknowledgements

We acknowledge the unconditional support by Director, CSIR-Central Institute for Medicinal and Aromatic Plants for providing Intuitional support through P-07 and P-50 grant. We thank the scientific community for developing and maintaining the open-source tools used in this study.

## Author Contributions

P.K.T, S.K., S.T. conceived the idea, S.T. designed the experiments, performed experiments and data analysis and wrote the manuscript. P.K.T, S.K., obtained funding to support this project. N.P.N. and S.V. maintained the plant materials. PKT, VK and SK provided supervision, guidance, and support., ST, SK, VG, and PKT revised and improved the manuscript.

